# Paxillin Promotes ATP-induced Activation of P2X7 Receptor and NLRP3 Inflammasome

**DOI:** 10.1101/2020.04.03.023721

**Authors:** Wenbiao Wang, Dingwen Hu, Yuqian Feng, Caifeng Wu, Yunting Song, Weiyong Liu, Aixin Li, Yingchong Wang, Keli Chen, Mingfu Tian, Feng Xiao, Qi Zhang, Weijie Chen, Pan Pan, Pin Wan, Yingle Liu, Kailang Wu, Jianguo Wu

## Abstract

The stimulation of P2X7 receptor by extracellular ATP leads to activation of NLRP3 inflammasome and release of pro-inflammatory cytokines. Here, we reveal a distinct mechanism by which Paxillin promotes ATP-induced activation of P2X7 receptor and NLRP3 inflammasome. Extracellular ATP induces Paxillin phosphorylation and facilitates Paxillin-NLRP3 interaction. Interestingly, Paxillin enhances NLRP3 deubiquitination and activates NLRP3 inflammasome upon ATP treatment and K^+^ efflux. Moreover, we reveal that UPS13 is a key enzyme for Paxillin-mediated NLRP3 deubiquitination upon ATP treatment. Notably, extracellular ATP promotes Paxillin and NLRP3 migration from cytosol to plasma membrane and facilitates P2X7-Paxillin interaction and Paxillin-NLRP3 association, resulting in the formation of P2X7-Paxillin-NLRP3 complex. Functionally, Paxillin is essential for ATP-induced NLRP3 inflammasome activation in mouse BMDMs and BMDCs as we as in human PBMCs and THP-1-differentiated macrophages. Thus, Paxillin plays key roles in ATP-induced activation of P2X7 receptor and NLRP3 inflammasome by facilitating the formation of the P2X7-Paxillin-NLRP3 complex.

## Introduction

The NACHT, LRR and PYD domains-containing protein 3 (NALP3), one of the host pattern recognition receptors (PRRs), recognizes pathogen-associated molecular patterns (PAMPs) and danger associated molecular patterns (DAMPs). NLRP3 controls maturation and secretion of interleukin-1β (IL-1β) and IL-18, two pleiotropic cytokines playing crucial roles in innate immune and inflammatory responses as well as instructing adaptive immune responses (Schroder and Tschopp, 2010). NLRP3 (the cytoplasmic sensor molecular) together with apoptosis-associated speck-like protein with CARD domain (ASC) (the adaptor protein) promote the cleavage of the pro-Caspase-1 (the effector protein) to generate active subunits p20 and p10, which regulate the maturation of IL-1β (Martinon et al., 2004). NLRP3 requires two signals for canonical activation and for IL-1β secretion: the first signal primes the cell to express NLRP3 and pro-IL-lβ mRNAs, and the second signal induces inflammasome assembly and activation (Martinon et al., 2009). NLRP3 forms a scaffold with ASC to provide a molecular platform for activation of pro-Caspase-1, which collectively comprises the inflammasome (Martinon et al., 2002). Activated Caspase-1 cleaves pro-IL-1β into active IL-1β, which is then secreted.

Extracellular adenosine triphosphate (ATP), a key DAMP, is released during inflammation by injured parenchymal cells, dying leukocytes, and activated platelets (Di Virgilio, 2007). Activation of NLRP3 inflammasome requires the activation of P2X7 receptor (P2X7R), a plasma membrane channel that is directly activated by extracellular ATP (Latz et al., 2013; Surprenant and North, 2009). After binding with ATP, the channel opens and induces transmembrane ion fluxes of K^+^, which is a key trigger of the NLRP3 inflammasome activation (Di Virgilio et al., 2017; Franceschini et al., 2015). However, the mechanism underlying this regulation is not fully understood and the molecule connecting P2X7 receptor and NLRP3 inflammasome is not identified.

Paxillin is a multi-domain protein that localizes to the extracellular matrix (ECM) and plays important roles in cell motility, adhesion, migration, and growth control (Christopher, 1998). The primary function of Paxillin is as a molecular adapter or scaffold protein that provides multiple docking sites at the plasma membrane for many proteins, such as Focal adhesion kinase (FAK) (Hanks et al., 2003) and extracellular signal-regulated kinase (ERK) (Ishibe et al., 2003). It is also involved in the efficient processing of Integrin- and growth factor-mediated signals derived from the extracellular environment (Nicholas and Christopher, 2008).

This study reveals a distinct mechanism underlying ATP-induced activation of P2X7 receptor and NLRP3 inflammasome mediated by Paxillin. Extracellular ATP induces Paxillin phosphorylation, facilitates Paxillin-NLRP3 interaction, and promotes NLRP3 deubiquitination, thereby activating NLRP3 inflammasome. Additionally, ATP promotes Paxillin and NLRP3 membrane migration and facilitates P2X7-Paxillin interaction, resulting in the formation of P2X7-Paxillin-NLRP3 complex. Moreover, ubiquitin proteasome system 13 (UPS13) is key enzyme for Paxillin-mediated NLRP3 deubiquitination (Liu et al., 2011). Paxillin functionally is essential for ATP- and Nigericin-induced NLRP3 inflammasome activation. Taken together, these results demonstrate that Paxillin promotes ATP-induced activation of P2X7 receptor and NLRP3 inflammasome by facilitating the formation of the P2X7-Paxillin-NLRP3 complex.

## Results

### Paxillin binds directly to LRR domain of NLRP3 in the cytosol

We have recently revealed that several proteins control the NLRP3 inflammasome activation through different mechanisms (Pan et al., 2019; Wan et al., 2019; Wang et al., 2017; Wang et al., 2018). Here, we further showed that Paxillin interacted with NLRP3 based on a yeast two-hybrid screen (Fig 1A). Co-immunoprecipitation (Co-IP) assays revealed that Paxillin interacted with NLRP3, but not with ASC or pro-Caspase-1 (Fig 1B). NLRP3 contains several prototypic domains, including a PYRIN domain, an NACHT domain, and seven LRR domains (Ye and Ting, 2008). Paxillin was co-precipitated with NLRP3, NACHT, and LRR, but not with PYRIN (Fig 1C). Glutathione S-transferase (GST) pull-down assays showed that GST-LRR was pulled down with Paxillin (Fig 1D) and GST-Paxillin was pulled down with NLRP3 (Fig 1E), suggesting that Paxillin directly binds to NLRP3 LRR domain. Confocal microscopy revealed that NLRP3 alone or Paxillin alone was diffusely distributed in the cytosol, whereas NLRP3 and Paxillin together were co-localized in the cytosol (Fig 1F). Collectively, these data demonstrate that Paxillin binds directly to LRR domain of NLRP3 in the cytosol.

**Figure 1.**
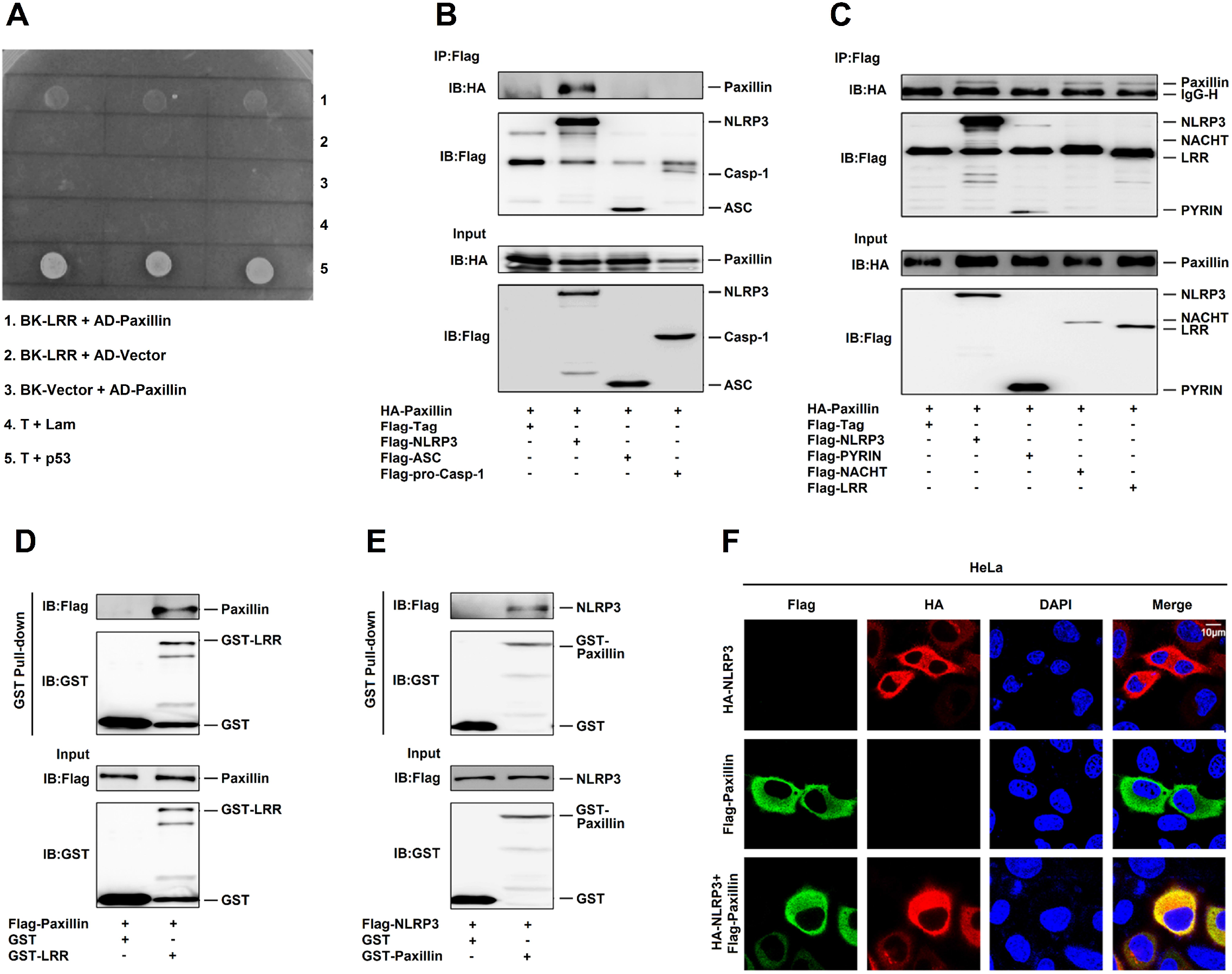
Paxillin binds directly to LRR domain of NLRP3 in the cytosol. **A** Identification of NLRP3 LRR domain-Paxillin interaction by yeast two-hybrid analysis. Yeast strain AH109 cells were transformed with the combination of BD and AD plasmid, as indicated. Transformed yeast cells were first grown on the SD-minus Trp/Leu plates for three days. The colony of yeast was then streaked on SD-minus Trp/Leu/Ade/His plates (QDO). BD-p53 and AD-T was used as a positive control and BD-Lam and AD-T as a negative control. **B** HEK293T cells were co-transfected with pHA-Paxillin and pFlag-Vector, pFlag-NLRP3, pFlag-ASC, and pFlag-Casp-1. Lysates were prepared and subjected to IP using anti-Flag antibody and analyzed by immunoblotting using an anti-HA or anti-Flag antibody (top) or subjected directly to Western blot using an anti-Flag or anti-HA antibody (as input) (bottom). **C** HEK293T cells were co-transfected with pHA-Paxillin and pFlag-Vector, pFlag-NLRP3, pFlag-PYRIN, pFlag-NACHT, and pFlag-LRR. Lysates were prepared and subjected to IP using anti-Flag antibody and analyzed by immunoblotting using an anti-HA or anti-Flag antibody (top) or subjected directly to Western blot using an anti-Flag or anti-HA antibody (as input) (bottom). **D** Extracts from HEK293T cells transfected with Flag-Paxillin were incubated with 10 μg GST proteins or GST-LRR protein that was incubated with glutathione-Sepharose beads. Mixture were washed three times and then analyzed by immunoblotting using anti-Flag and anti-GST antibody (top). Lysates from transfected HEK293T cells were analyzed by immunoblotting using anti-Flag antibody (as input) (bottom). **E** Extracts from HEK293T cells transfected with Flag-NLRP3 were incubated with 10 μg GST proteins or GST-Paxillin protein which was incubated with glutathione-Sepharose beads. Mixture were washed three times and then analyzed by immunoblotting using anti-Flag and anti-GST antibody (top). Lysates from transfected HEK293T cells were analyzed by immunoblotting using anti-Flag antibody (as input) (bottom). **F** HeLa cells were transfected with pFlag-Paxillin, pHA-NLRP3, or co-transfected with pFlag-Paxillin and pHA-NLRP3. Subcellular localization of Flag-Paxillin (green), HA-NLRP3 (red) and the nucleus marker DAPI (blue) were examined by confocal microscopy.

### ATP promotes Paxillin phosphorylation and Paxillin-NLRP3 interaction

ATP and Nigericin are common activators of the NLRP3 inflammasome (Mariathasan et al., 2006). Human acute monocytic leukemia cells (THP-1) that stably expressed Paxillin were generated by infection with Paxillin-Lentivirus, which were then treated with ATP or Nigericin. ATP-induced Inflammasome activation, as indicated by IL-1β secretion, IL-1β maturation, and Caspase-1 cleavage, was further promoted by Paxillin, whereas Nigericin-induced inflammasome activation was not affected by Paxillin (Fig 2A, B), suggesting that Paxillin specifically promotes ATP-induced NLRP3 inflammasome activation. As ATP stimulates P2X7 ATP-gated ion channel and triggers K^+^ efflux, leading to inflammasome activation (Kahlenberg and Dubyak, 2004), we evaluated the effect of Paxillin on NLRP3 inflammasome activation. In Paxillin stable THP-1 cells, IL-1β secretion, IL-1β maturation, and Caspase-1 cleavage were induced by ATP and further facilitated by Paxillin in the presence of Dimethylsulphoxide (DMSO) or BAPTA-AM (Ca^2+^ chelator), whereas activations of IL-1β and Caspase-1 were not induced by ATP or Paxillin in the presence of YVAD (Caspase-1 inhibitor), Glybenclamide (K^+^ efflux inhibitor), A438079 (P2X7 antagonist), or AZ10606120 (P2X7 antagonist) (Fig 2C, D). These results indicate that like ATP, Paxillin promotes IL-1β and Caspase-1 activation depending on P2X7, K^+^ efflux, and Casp-1, and suggest that Paxillin facilitates ATP-induced NLRP3 inflammasome activation.

**Figure 2.**
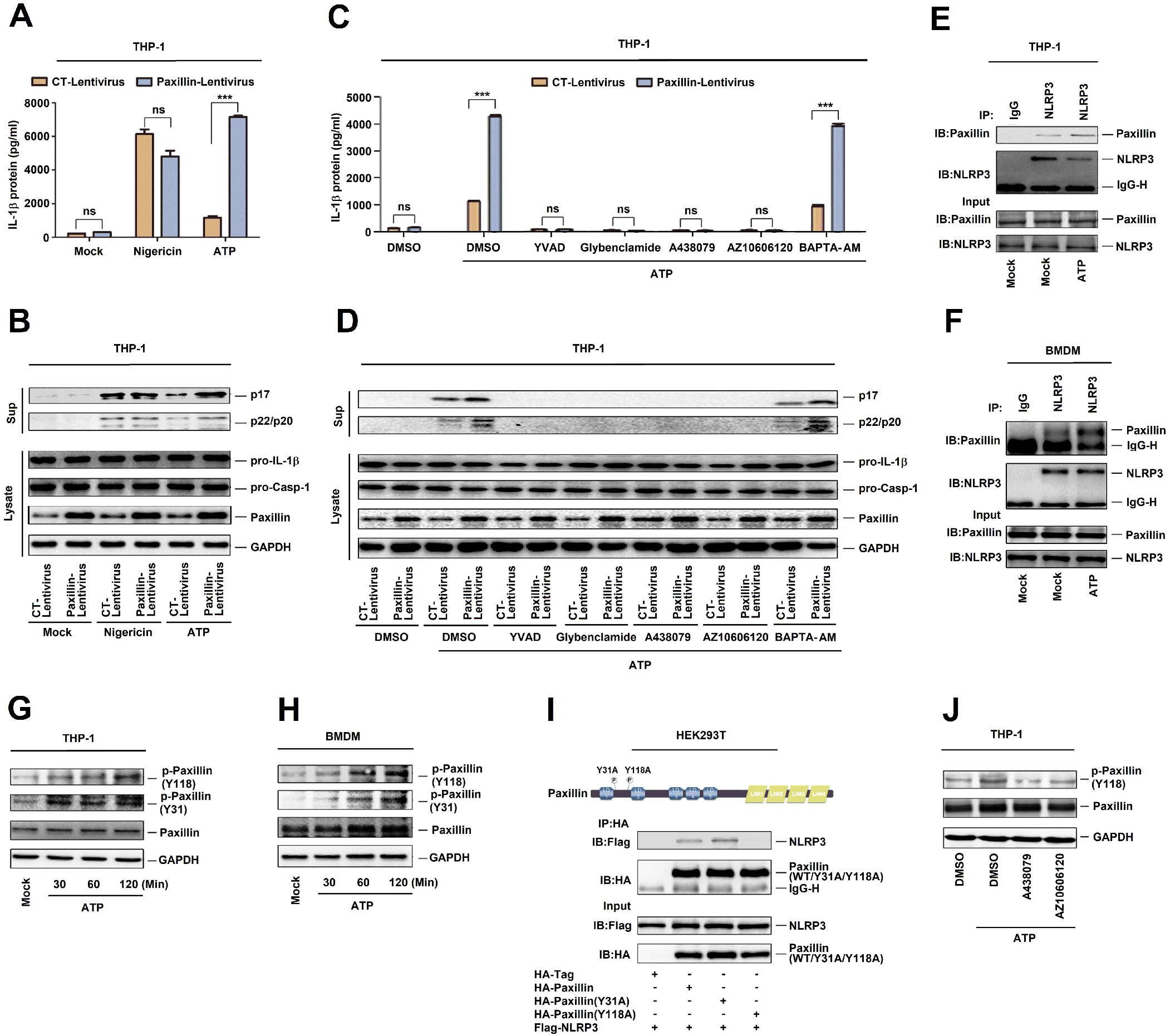
ATP promotes Paxillin phosphorylation and Paxillin-NLRP3 interaction. **A–D** THP-1 cells stably expressing control lentivirus or Paxillin lentivirus were generated and differentiated to macrophages, which were then treated with ATP (5 mM) or Nigericin (2 μM) for 2 h (A, B), treated with DMSO, YVAD (50 μM), Glybenclamide (25 μg/ml), A438079 (100 μM), AZ10606120 (100 μM), and BAPTA-AM (30 μM) for 1 h before the treatment with ATP (5 mM) for 2 h (C, D). IL-1β levels in supernatants were determined by ELISA (A, C). Mature IL-1β (p17) and cleaved Casp-1 (p22/p20) in supernatants or Paxillin, pro-IL-1β, and pro-Casp-1 in lysates were determined by Western blot (B, D). **E** TPA-differentiated THP-1 macrophages were treated with DMSO or ATP (5 mM) for 2 h. Lysates were prepared and subjected to IP (top) or subjected to Western blot (as input) (bottom). **F** BMDM cells prepared from C57BL/6 mice bone marrow were treated with LPS (1 μg/ml) for 6 h, and the primed BMDMs were stimulated by DMSO or ATP (5 mM) for 30 min. Lysates were prepared and subjected to IP (top) or subjected to Western blot (as input) (bottom). **G, H** TPA-differentiated THP-1 macrophages were treated with ATP (5 mM) for 30, 60, and 120 min (G). LPS-primed BMDMs were stimulated by ATP (5 mM) for 5, 15, and 30 min (H). The protein level of p-Paxillin(Y 118), p-Paxillin(Y31), Paxillin, and GAPDH was determined by Western blot. **I** HEK293T cells were co-transfected with pFlag-NLRP3 and pHA-Vector, pHA-Paxillin, pHA-Paxillin-(Y31A), and pHA-Paxillin-(Y118A). Lysates were prepared and subjected to IP (top) or subjected to Western blot (as input) (bottom). **J** TPA-differentiated THP-1 macrophages were treated with DMSO, A438079 (100 μM), AZ10606120 (100 μM) for 1 h before the treatment with ATP (5 mM) for 2 h. The protein level of p-Paxillin(Y118), Paxillin, and GAPDH was determined by Western blot. Data shown are means ± SEMs; ***p < 0.0001. ns, no significance.

The mechanism by which Paxillin regulates ATP-induced NLRP3 inflammasome was investigated. As Paxillin interacts with NLRP3, we explored whether ATP affects this interaction. Paxillin-NLRP3 interaction was enhanced by ATP in TPA-differentiated THP-1 macrophages (Fig 2E), and LPS-primed mouse bone marrow derived macrophages (BMDMs) (Fig 2F). Paxillin is a phosphorylated molecule and plays an important role in cell adhesion and migration (Schaller, 2001), and phosphorylations of Paxillin at Y31 and Y118 were essential for the generation of protein binding module (Hanks et al., 2003). We revealed that Paxillin Y31 and Y118 phosphorylations were promoted by ATP in THP-1 differentiated macrophages (Fig 2G) and LPS-primed BMDMs (Fig 2H). Like Paxillin, Paxillin(Y31A) interacted with NLRP3, whereas Paxillin(Y118A) failed to interact with NLRP3 (Fig 2I), suggesting an essential role of Y118 in Paxillin-NLRP3 interaction. Moreover, ATP-induced Paxillin Y118 phosphorylation was attenuated by P2X7 antagonists (A438079 and AZ10606120) (Fig 2J), indicating that P2X7 is involved in Paxillin Y118 phosphorylation. Therefore, ATP- and P2X7-induced Paxillin Y118 phosphorylation is essential for Paxillin-NLRP3 interaction.

### ATP stimulates P2X7-Paxillin interaction

Extracellular ATP stimulates P2X7 ATP-gated ion channel and inducing gradual recruitment of Pannexin-1 membrane pore (Kanneganti et al., 2007). As the downstream adaptor of Integrin, Paxillin functions as a scaffold for the recruitment of proteins into a complex (Franceschini et al., 2015), we thus explored whether Paxillin interacts with P2X7. The result revealed that Paxillin interacted with P2X7 and Pannexin-1 (Fig 3A), and P2X7-Paxillin interaction was promoted by ATP (Fig 3B, C). Like Paxillin, Paxillin(Y31A) and Paxillin(Y118A) interacted with P2X7 (Fig 3D). Notably, P2X7-Paxillin(Y31A) interaction was not facilitated by ATP (Fig 3E), whereas P2X7-Paxillin(Y118A) interaction was promoted by ATP (Fig 3F). These results reveal that phosphorylation site Y31 is involved in ATP-induced P2X7-Paxillin interaction.

**Figure 3.**
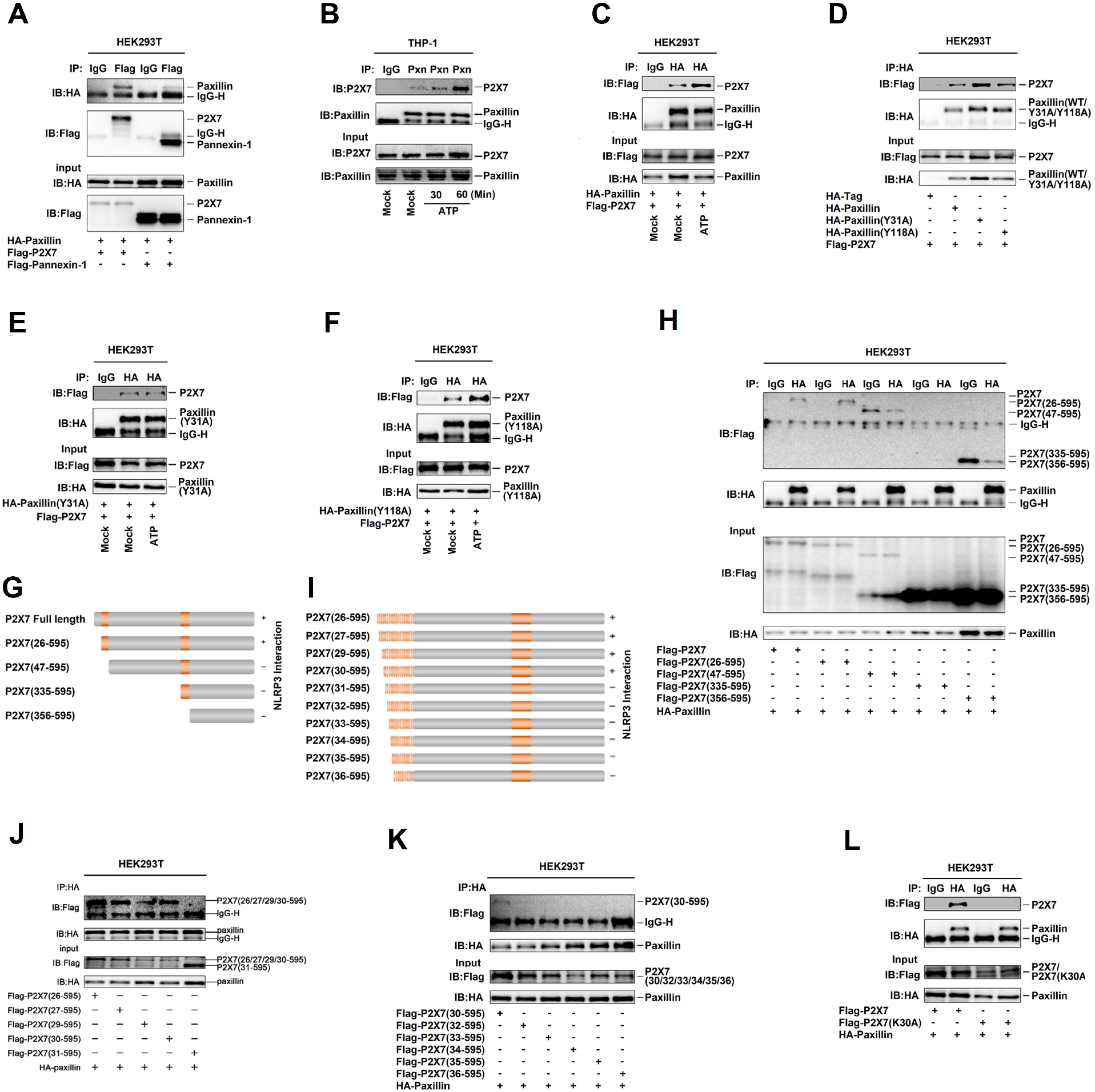
ATP stimulates P2X7-Paxillin interaction. **A** HEK293T cells were co-transfected with pFlag-P2X7 and pHA-Paxillin or pFlag-Pannexin-1 and pHA-Paxillin. **B** TPA-differentiated THP-1 macrophages were mock-treated or treated with ATP (5 mM) for 30 and 60 min. **C** HEK293T cells were co-transfected with pFlag-P2X7 and pHA-Paxillin, and treated with DMSO or ATP (5 mM) for 2 h. **D** HEK293T cells were co-transfected with pFlag-P2X7 and pHA-vector, pHA-Paxillin, pHA-Paxillin(Y31A), and pHA-Paxillin(Y118A). **E** HEK293T cells were co-transfected with pFlag-P2X7 and pHA-paxillin(Y31A), and treated with DMSO or ATP (5 mM) for 2 h. **F** HEK293T cells were co-transfected with pFlag-P2X7 and pHA-Paxillin(Y118A), and treated with DMSO or ATP (5 mM) for 2 h. **G** The diagrams of P2X7, P2X7(26–595), P2X7(47–595), P2X7(335–595), and P2X7(356–595) (left). **H** HEK293T cells were co-transfected with pHA-Paxillin and pFlag-P2X7, pFlag-P2X7(26–595), pFlag-P2X7(47–595), pFlag-P2X7(335–595), and pFlag-P2X7(356–595). **I** The diagrams of P2X7(26/27/29/30/31/32/33/34/35/36-595). **J** HEK293T cells were co-transfected with pHA-Paxillin and pFlag-P2X7(26–595) pFlag-P2X7(27–595), pFlag-P2X7(29–595), pFlag-P2X7(30–595) and pFlag-P2X7(31–595). **K** HEK293T cells were co-transfected with pHA-Paxillin and pFlag-P2X7(30–595) pFlag-P2X7(32–595), pFlag-P2X7(33–595), pFlag-P2X7(34–595), pFlag-P2X7(35–595) and pFlag-P2X7(36–595). **L** HEK293T cells were co-transfected with pFlag-P2X7 and pHA-Paxillin or pFlag-P2X7(K30A) and pHA-Paxillin.

P2X7 subunit contains intracellular N-termini (1–25aa), C-termini (357–595aa), and two transmembrane domains (TM1 and TM2) (47–334) (Schaller, 2001). Accordingly, five plasmids expressing P2X7 and deletion mutants were constructed (Fig 3G). Paxillin interacted withP2X7 and P2X7(26–595), whereas Paxillin failed to interact with P2X7(47–595), P2X7(335–595), or P2X7(356–595), suggesting that Paxillin interacts with 26aa–46aa of P2X7 (Fig 3H). To narrow down the interaction region, ten plasmids expressing truncated P2X7 proteins were constructed (Fig 3I). Paxillin interacted with P2X7(26–595), P2X7(27–595), P2X7(29–595), and P2X7(30–595), but it failed to associate with P2X7(31–595), P2X7(32–595), P2X7(33–595), P2X7(34–595), P2X7(35–595), or P2X7(36–595) (Fig 3J, K), indicating that P2X7 K30 is required for Paxillin interaction. To confirm this result, K30 was mutated to A30 to generate mutant P2X7(K30A). Notably, unlike P2X7, P2X7(K30A) failed to interact with Paxillin (Fig 3L), confirming that K30 is essential for the interaction with Paxillin. Therefore, ATP stimulates P2X7-Paxillin interaction, and promotes Paxillin-NLRP3 interaction, and thus Paxillin plays an essential role in P2X7-Paxillin-NLRP3 complex assembly upon ATP induction.

### Paxillin promotes NLRP3 deubiquitination depending on extracellular ATP and K^+^ efflux

As NLRP3 deubiquitination is important for the inflammasome activation (Py et al., 2013), here, the roles of ATP and Paxillin in NLRP3 deubiquitination were evaluated. The level of NLRP3 ubiquitination was attenuated by ATP in THP-1 differentiated macrophages, and LPS-primed BMDMs (Fig 4A, B). Similarly, the abundance of NLRP3 ubiquitination was reduced by Paxillin in HEK293T cells and HeLa cells (Fig 4C, D). In addition, ATP-induced NLRP3 deubiquitination was detected in the absence of sh-Paxillin, whereas it was repressed by sh-Paxillin in THP-1 cells and BMDMs (Fig 4E, F). Collectively, these results reveal that Paxillin promotes ATP-induced NLRP3 deubiquitination.

**Figure 4.**
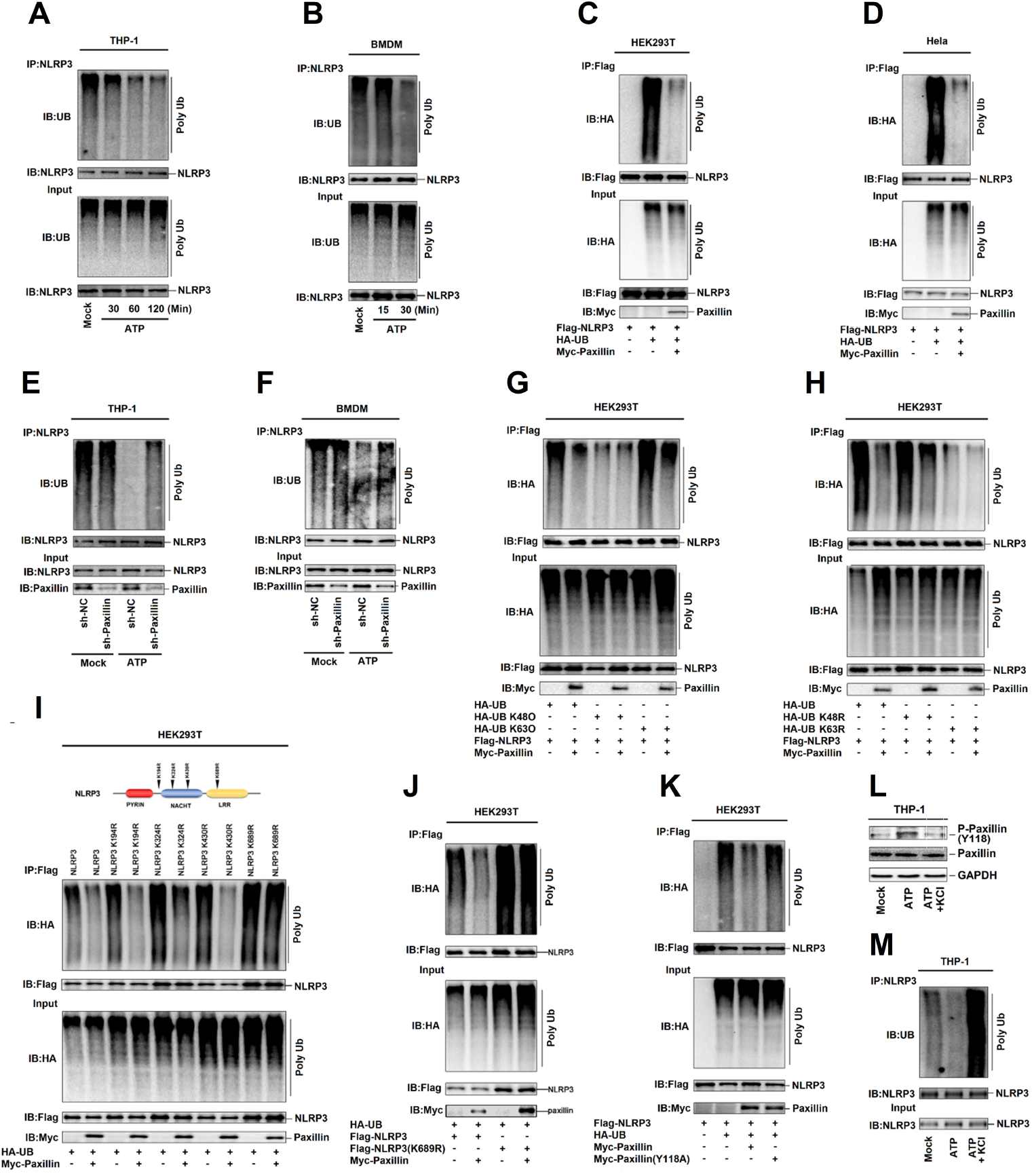
Paxillin promotes NLRP3 deubiquitination depending on extracellular ATP and K^+^ efflux. **A** TPA-differentiated THP-1 macrophages were treated with ATP (5 mM) for 30, 60, and 120 min. **B** BMDM cells prepared from C57BL/6 mice bone marrow were treated with LPS (1 μg/ml) for 6 h, and the primed BMDM cells were stimulated by ATP (5 mM) for 15 and 30 min. **C, D** HEK293T cells (C) and Hela cells (D) were co-transfected with pFlag-NLRP3, pHA-Ubiquitin, or pMyc-Paxillin. **E** THP-1 cells stably expressing shRNA targeting Paxillin was generated and differentiated to macrophages, which were then treated with ATP (5 mM) for 2 h. **F** BMDMs prepared from C57BL/6 mice bone marrow were infected by lentivirus that express shRNA targeting Paxillin for 3 days. Before stimulation, the BMDMs were treated with LPS (1 μg/ml) for 6 h, and the primed BMDMs were stimulated by ATP (5 mM) for 30 min. **G** HEK293T cells were co-transfected with pFlag-NLRP3, pHA-UB, pHA-UB(K48O), pHA-UB(K63O) or pMyc-Paxillin. **H** HEK293T cells were co-transfected with pFlag-NLRP3, pHA-UB, pHA-UB(K48R), pHA-UB(K63R), or pMyc-Paxillin. **I** HEK293T cells were co-transfected with pFlag-NLRP3, pFlag-NLRP3(K194R), pFlag-NLRP3(K324R), pFlag-NLRP3(K403R), pFlag-NLRP3(K689R), pHA-UB, or pMyc-Paxillin. **J** HEK293T cells were co-transfected with pFlag-NLRP3, pFlag-NLRP3(K689R), pHA-UB, or pMyc-Paxillin. **K** HEK293T cells were co-transfected with pFlag-NLRP3, pHA-UB, pMyc-Paxillin, or pMyc-Paxillin(Y 118A). **L** TPA-differentiated THP-1 macrophages were treated with ATP (5 mM) for 2 h and with/without 50 mM extracellular KCl. The protein level of p-Paxillin-(y118), Paxillin and GAPDH was determined by Western blot. **M** TPA-differentiated THP-1 macrophages were treated with ATP (5 mM) for 2 h in the presence or absence of 50 mM extracellular KCl. Lysates were prepared and subjected to denature-IP (top) or subjected to Western blot (as input) (bottom) (A–M).

To determine the nature of NLRP3 ubiquitination, two ubiquitin mutants that retains only a single lysine residue (KO) and two ubiquitin mutants in which only one lysine residue is mutated (KR) were generated. NLRP3 was ubiquitinated by UB, UB(K63O), and UB(K48R), whereas it failed to be ubiquitinated by UB(K48O) or UB(K63R), and NLRP3 K63-linked ubiquitination was attenuated by Paxillin (Fig 4G, H), indicating that NLRP3 ubiquitination is K63-linked and Paxillin facilitates the removal of NLRP3 K63-linked ubiquitination.

Moreover, the site of NLPR3 ubiquitination was determined by constructing and analyzing four only one lysine residual mutants (KR) of NLPR3 (Fig 4I, top). Ubiquitinations of NLRP3, NLRP3(K194R), and NLRP3(K430R) were attenuated by Paxillin, ubiquitination of NLRP3(K324R) was down-regulated by Paxillin, whereas ubiquitination of NLRP3(K689R) was not affected by Paxillin (Fig 4I, bottom), and Paxillin attenuated NLRP3 ubiquitination but failed to reduce NLRP3(K689R) ubiquitination (Fig 4J), revealing that Paxillin specifically removes NLRP3 ubiquitination at K689. As Paxillin Y118 phosphorylation is required for Paxillin-NLRP3 interaction, we evaluated its effect on NLRP3 deubiquitination. NLRP3 ubiquitination was notably attenuated by Paxillin, whereas it was not affected by Paxillin(Y118A) (Fig 4K), indicating that Paxillin Y118 phosphorylation is required for NLRP3 deubiquitination.

As ATP-induced Paxillin Y118 phosphorylation is essential for Paxillin-NLRP3 interaction and deubiquitination, we further explored the effect of K^+^ efflux on Paxillin Y118 phosphorylation. Paxillin Y118 phosphorylation was induced by ATP, but such induction was attenuated by KCl treatment (Fig 4L). Furthermore, the role of K^+^ efflux in NLRP3 deubiquitination was revealed. Notably, ATP attenuated NLRP3 ubiquitination, but such attenuation was suppressed by KCl (Fig 4M), demonstrating that K^+^ efflux is important for Paxillin- and ATP-induced deubiquitination of NLRP3. Together, we reveal that Paxillin facilitates the remove NLRP3 K63-linked ubiquitination at K689 and that ATP and K^+^ efflux are critical for Paxillin-mediated NLRP3 deubiquitination.

### UPS13 is essential for Paxillin-mediated NLRP3 deubiquitination upon ATP treatment

Next, the enzyme required for Paxillin-mediated NLRP3 deubiquitination was then determined. As deubiquitinating enzymes, such as eukaryotic translation initiation factor 3, subunit 5 (EIF3S5), ubiquitin proteasome system 13 (UPS13), and OTU domain-containing ubiquitin aldehyde-binding protein 1 (OTUB1), were potentially involved in the deubiquitination of NLRP3 (Py et al., 2013) [25]. Here, we explored the roles of these enzymes in NLRP3 deubiquitination. Paxillin interacted with EIF3S5, UPS13 (Fig 5A), revealing that EIF3S5 and UPS13 might be participated in Paxillin-mediated NLRP3 deubiquitination. To confirm the roles of EIF3S5 and UPS13 in the regulation of NLRP3 activation, Hela cells stably expressing sh-RNA targeting human EIF3S5 (sh-EIF3S5) and human UPS13 (sh-UPS13) were generated and analyzed (Fig 5B). NLRP3 ubiquitination was attenuated by Paxillin in the presence of sh-NC and sh-EIF3S5, whereas it was relatively unaffected by Paxillin in the presence of sh-UPS13 (Fig 5C), indicating that UPS13 is required for Paxillin-mediated NLRP3 deubiquitination. Moreover, co-IP assays revealed that Paxillin and UPS13 interacted with each other (Fig 5D–G). Like NLRP3, NLRP3 NACHT domain and LRR domain interacted with UPS13, but NLRP3 PYRIN domain failed to interact with UPS13 (Fig 5H).

**Figure 5.**
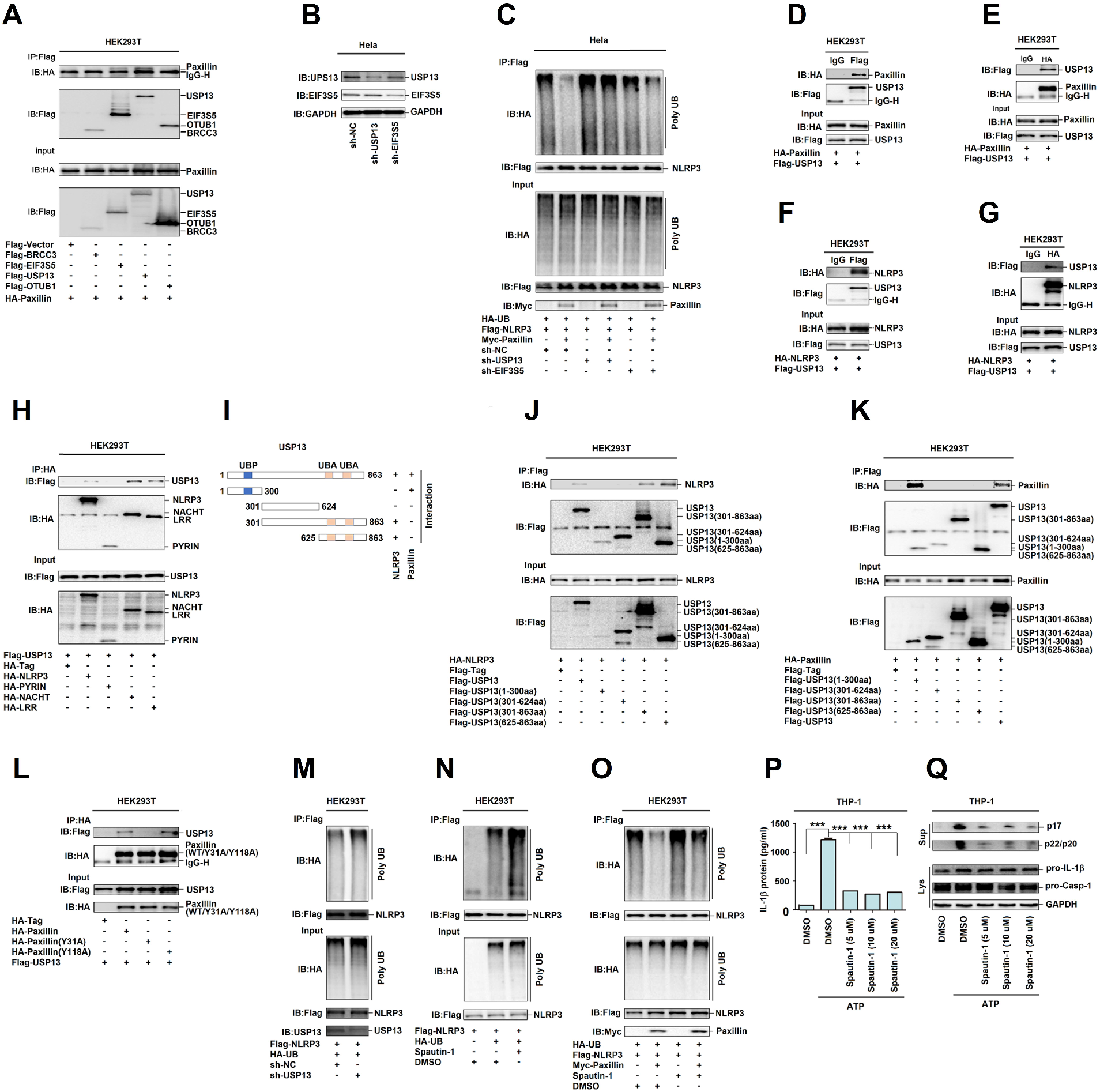
UPS13 is essential for Paxillin-mediated NLRP3 deubiquitination upon ATP treatment. **A** HEK293T cells were co-transfected with pHA-Paxillin and pFlag-BRCC3, pFlag-EIF3S5, pFlag-UPS13 or pFlag-OTUB1. Lysates were prepared and subjected to IP (top) or subjected to Western blot (as input) (bottom). **B** Hela cells stably expressing sh-UPS13 or sh-EIF3S5 were generated and analyzed. **C** The stable Hela cells were co-transfected with pFlag-NLRP3, pHA-UB, or pMyc-Paxillin. Lysates were prepared and subjected to denature-IP (top) or subjected to Western blot (bottom). **D, E** HEK293T cells were transfected with pFlag-UPS13 and pHA-Paxillin. **F, G** HEK293T cells were transfected with pFlag-UPS13 and pHA-NLRP3. Lysates were prepared and subjected to IP (top) or subjected to Western blot (bottom) (D–G). **H** HEK293T cells were co-transfected with pFlag-UPS13 and pHA-NLRP3 or pHA-NLRP3 mutants. **I** The diagrams of UPS13 and its mutants, as indicated. **J** HEK293T cells were co-transfected with pHA-NLRP3 and pFlag-UPS13 or pFlag-UPS13 mutants. **K** HEK293T cells were co-transfected with pHA-Paxillin and pFlag-UPS13 or pFlag-UPS13 mutants. **L** HEK293T cells were co-transfected with pFlag-UPS13 and pHA-Paxillin or pHA-Paxillin mutants. Lysates were prepared and subjected to IP (top) or subjected to Western blot (bottom) (H, j –l). **M** HEK293T cells stably expressing sh-UPS13 were generated. The stable cells were co-transfected with pFlag-NLRP3 and pHA-UB. **N** The stable HEK293T cells were co-transfected with pFlag-NLRP3, or pHA-UB with or without Spautin-1 (20 μM) for 6 h. **O** The stable HEK293T cells wre co-transfected with pFlag-NLRP3, pHA-UB, or pMyc-Paxillin with or without Spautin-1 (20 μM) for 6 h. Lysates were prepared and subjected to denature-IP (top) or subjected to Western blot (bottom) (m–o). **P**, **Q** TPA-differentiated THP-1 macrophages were treated with ATP (5 mM) for 2 h with/without Spautin-1 (5, 10, and 20 μM). IL-1β in supernatants was determined by ELISA (P). Mature IL-1β(p17) and cleaved Caspase-1(p22/p20) in supernatants and Paxillin, pro-IL-1β, and pro-Casp-1 in lysates were determined by Western blot (Q). Data shown are means ± SEMs; ***p < 0.0001.

In another hand, the domains of UPS13 required for UPS13-NLRP3 interaction and UPS13-Paxillin association were assessed by constructing and analyzing plasmids encoding for WT UPS13(1–863) and four truncated proteins (Fig 5I). Like WT UPS13(1–863), UPS13(301–863) and UPS13(625–863) interacted with NLRP3, but UPS13(1–300) or UPS13(301–624) failed to interact with NLRP3 (Fig 5J), indicating that 625aa–863aa of UPS13 are involved in UPS13–NLRP3 interaction. Additionally, UPS13(1–863) and UPS13(1–300) interacted with Paxillin, but UPS13(301–624), UPS13(301–863), and UPS13(625–863) could not interact with Paxillin (Fig 5K), indicating that 1aa-300aa of UPS13 is involved in UPS13-Paxillin interaction. Furthermore, Paxillin and Paxillin(Y118A) interacted with UPS13, but Paxillin(Y31A) failed to associate with UPS13 (Fig 5L), implicating that Paxillin Y31 phosphorylation is important for UPS13-Paxillin interaction.

To determine the role of UPS13 in the regulation of NLRP3 ubiquitination, we generated HEK293T cells stably expressing sh-UPS13. The level of NLRP3 ubiquitination was promoted by sh-UPS13 (Fig 5M), facilitated by Spautin-1 (an inhibitor of deubiquitinating enzyme activity of UPS13) (Fig 5N), attenuated by Paxillin, whereas Paxillin-mediated attenuation of NLRP3 ubiquitination was repressed by Spautin-1 (Fig 5O), indicating that UPS13 is required for NLRP3 deubiquitination. Moreover, IL-1β secretion, IL-1β maturation, and Casp-1 cleavage induced by ATP were repressed by Spautin-1 (Fig 5P, Q), suggesting that UPS13 deubiquitinating enzyme activity is critical for ATP-induced NLRP3 inflammasome activation. Taken together, these data reveal that UPS13 is essential for Paxillin-mediated deubiquitination of NLRP3 upon ATP treatment.

### ATP induces Paxillin and NLRP3 membrane migration to facilitate P2X7-Paxillin-NLRP3 complex formation

As Paxillin is a scaffold for the recruitment of proteins into a complex apposing to the plasma membrane (Martinon et al., 2004), and Paxillin facilitates the P2X7-Paxillin-NLRP3 complex assembly, we thus explored whether Paxillin recruits NLRP3 to plasma membrane. In Hela cells, NLRP3 alone and Paxillin alone diffusely distributed in the cytosol in the absence of ATP; NLRP3 forms small spots in the cytosol and Paxillin localized in the membrane, whereas NLRP3 and Paxillin together co-localized and formed membrane blebbing in the presence of ATP (Fig 6A), similar to that described previously (Pelegrin and Surprenant, 2006). In TPA-differentiated THP-1 macrophages, endogenous NLRP3 was diffusely distributed in the cytosol without ATP but formed spots in the membrane as indicated by Dil (red) (a dye of cell membrane) in the presence of ATP (Fig 6B); endogenous phosphorylated Paxillin was hardly detected without ATP, whereas it was detected in the membrane upon ATP treatment (Fig 6C); and notably, endogenous NLRP3 and phosphorylated Paxillin co-localized and formed spots in the membrane upon ATP treatment (Fig 6D). Moreover, in LPS-primed BMDMs, endogenous NLRP3 was diffusely distributed in the cytosol without ATP, but localized and formed spots in the membrane upon ATP treatment (Fig 6E); endogenous phosphorylated Paxillin was hardly detected without ATP, but localized in the membrane after ATP treatment (Fig 6F); and endogenous NLRP3 and phosphorylated Paxillin co-localized and formed spots in the membrane after treated with ATP (Fig 6G). Collectively, these results suggest that ATP induces NLRP3 and Paxillin translocation from cytosol to plasma membrane.

**Figure 6.**
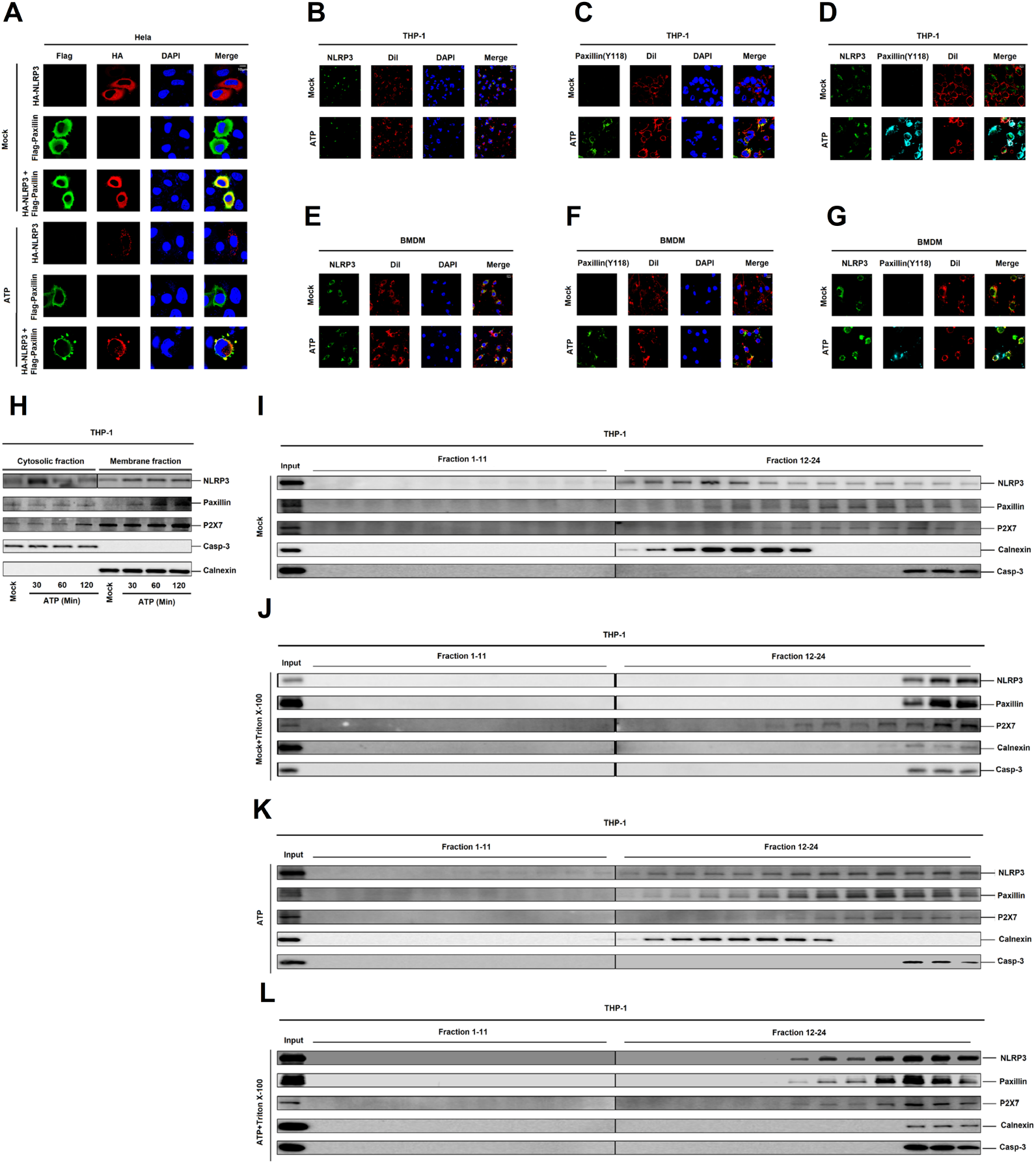
ATP induces Paxillin and NLRP3 membrane migration to facilitate the formation of P2X7-Paxillin-NLRP3 complex. **A** Hela cells were transfected with pFlag-Paxillin, pHA-NLRP3, or co-transfected with pFlag-Paxillin and pHA-NLRP3, and treated with ATP (5 mM) for 2 h. Subcellular localization of Flag-Paxillin (green), HA-NLRP3 (red), and the nucleus marker DAPI (blue) were examined by confocal microscopy. **B** TPA-differentiated THP-1 macrophages were treated with ATP (5 mM) for 2 h. Subcellular localization of NLRP3 (green), the membrane marker Dil (red), and DAPI were examined by confocal microscopy. **C** TPA-differentiated THP-1 macrophages were treated with ATP (5 mM) for 2 h. Subcellular localization of Paxillin(Y118) (green), Di l(red), and DAPI (blue) were examined by confocal microscopy. **D** TPA-differentiated THP-1 macrophages were treated with ATP (5 mM) for 2 h. Subcellular localization of NLRP3 (green), Paxillin(Y118) (cyan), and Dil (red) were examined by confocal microscopy. **E** LPS-primed BMDMs were stimulated by ATP (5 mM) for 30 min. Subcellular localization of NLRP3 (green), Dil (red), and DAPI (blue) were examined by confocal microscopy. **F** LPS-primed BMDMs were stimulated by ATP (5 mM) for 30 min. Subcellular localization of Paxillin(Y118) (green), Dil (red), and DAPI (blue) were examined by confocal microscopy. **G** LPS-primed BMDMs were stimulated by ATP (5 mM) for 30 min. Subcellular localization of NLRP3 (green), Paxillin(Y118) (cyan), and Dil (red) were examined by confocal microscopy. **H** Subcellular fractionation of TPA-differentiated THP-1 macrophages in the presence of ATP (5 mM) for 30, 60, and 120 min. The protein levels of NLRP3, Paxillin, P2X7, Caspase-3, and Calnexin in cytosolic fraction (left) and membrane fraction (right) were determined by Western blot. **I–L** Membrane floatation assays of TPA-differentiated THP-1 macrophages post-nuclear lysates on a 10–45% Optiprep gradient under the treatment of ATP (5 mM) for 30 min in the presence or absence of 1% Triton X-100. The protein level of NLRP3, Paxillin, P2X7, Caspase-3, and Calnexin was determined by Western blot in each 24 fractions and used as the input.

The distribution of NLRP3 in TPA-differentiated THP-1 macrophages upon ATP treatment was examined. Fractionation fidelity was verified by Caspase-3 in the cytosolic fraction and Calnexin in the membrane fraction. Without ATP treatment, NLRP3, Paxillin, and P2X7 localized in the cytosolic fraction and membrane fraction, Caspase-3 localized in the cytosolic fraction, and Calnexin distributed in membrane fraction; upon ATP treatment, NLRP3, Paxillin, and P2X7 distributed in the membrane fraction (Fig 6H), suggesting that NLRP3 and Paxillin migrate from the cytosol to membrane upon ATP stimulation. Membrane flotation assay can indicate that membrane-bound organelles migrate from dense to light fractions (Barnett et al., 2019), and membrane-bound organelles are sensitive to Triton X-100 except plasma membrane (Brown and Rose, 1992; Lingwood and Simons, 2010). We thus performed membrane flotation assays on Optiprep gradients bottom-loaded with lysate supernatants of TPA-differentiated THP-1 macrophages. Without treatments, NLRP3 floated from the bottom fractions (21-24) into the membrane fractions (12-20), Paxillin floated from the bottom fractions (21 –24) into the membrane fractions (14–20), and P2X7 floated from the bottom fractions (21–24) into the lighter fractions (15–20), while Calnexin remained in the lighter membrane fraction (12–18) and Caspase-3 stayed in the bottom fractions (22–24) (Fig 6I). Notably, after Triton X-100 treatment, NLRP3, Paxillin, Calnexin, and Caspase-3 remained in the bottom fractions (22–24), whereas P2X7 floated from the bottom fractions (22–24) into the lighter fractions (17–21) (Fig 6J). Notably, upon ATP stimulation, NLRP3 floated from the bottom fractions (21 –24) into the membrane fractions (12–20), Paxillin floated from the bottom fractions (21–24) in to the lighter fractions (13–20), and P2X7 floated from the bottom fractions (21–24) into the membrane fractions (17–20); while Calnexin remained in the membrane fraction (12–19) and Caspase-3 retained in the bottom fractions (22–24) (Fig 6K). Interestingly, upon ATP stimulation and Triton X-100 treatment, NLRP3, Paxillin, and P2X7 floated from the bottom fractions (21–24) into the membrane fractions (18–20), Calnexin retained the bottom fractions (21–24), and Caspase-3 remained in the bottom fractions (22–24) (Fig 6L). Together, the results demonstrate that NLRP3, Paxillin, and P2X7 colocalized on plasma membrane upon ATP treatment, and suggest that ATP induces Paxillin and NLRP3 membrane migration to facilitate P2X7-Paxillin-NLRP3 complex formation.

### Paxillin is required for ATP- and Nigericin-induced NLRP3 inflammasome activation

At least three NLRP family members (NLRP1, NLRP3, and NLRC4/IPAF) and one HIN-200 family member (Absent in Melanoma 2, AIM2) have been reported to exhibit inflammasome activity (Schroder and Tschopp, 2010). The NLRP1 inflammasome is activated by mycobacterial DNA-binding protein (MDP) (Faustin et al., 2007). The NLRP3 inflammasome is induced by ATP (Mariathasan et al., 2006), monosodium urate (MSU) (Martinon et al., 2006), Nigericin (Mariathasan et al., 2006), and Alum (Al) (Hornung et al., 2008). The NLRC4 inflammasome is stimulated by bacteria (Franchi et al., 2006). AIM2 senses cytosolic double-stranded DNA (dsDNA) (Hornung et al., 2009). Here, we determined the biological roles of Paxillin in the regulation of the four inflammasomes. BMDMs stably expressed sh-Paxillin were generated by infected with lentivirus expressing sh-Paxillin. The stable BMDMs were treated with four inflammasome activators, MDP for NLRP1 inflammasome, ATP for NLRP3 inflammasome, dA:dT for AIM2 inflammasome, and Salmonella for NLRC4 inflammasome. ATP-induced IL-1β secretion, IL-1β maturation, and Caspase-1 cleavage were attenuated by sh-Paxillin, whereas MDP-, dA:dT-, or Salmonella-induced IL-1β secretion, IL-1β maturation, and Caspase-1 cleavage were not affected by sh-Paxillin (Fig 7A, B), demonstrating that Paxillin is essential for the NLRP3 inflammasome activation.

**Figure 7.**
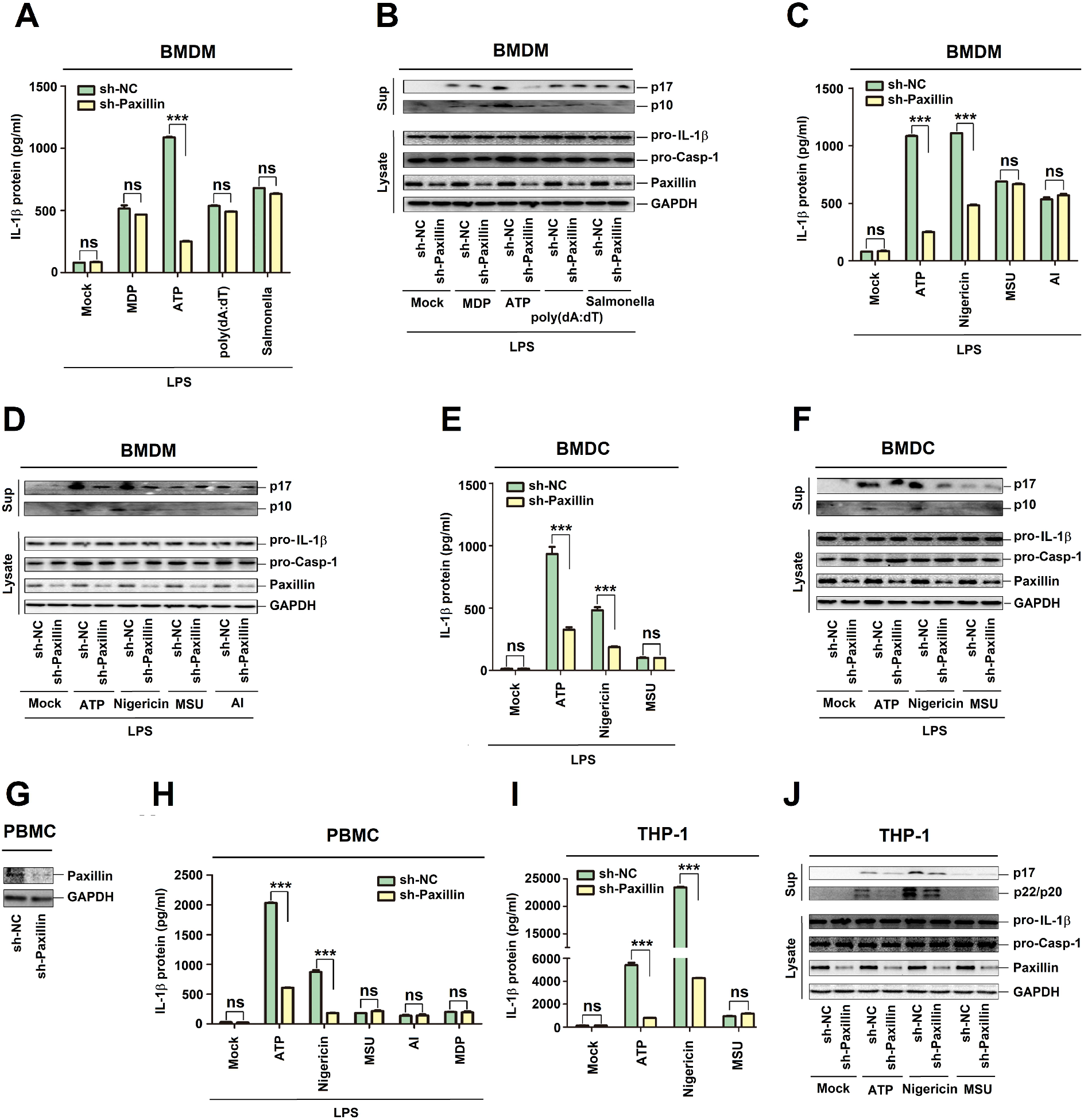
Paxillin is required for ATP- and Nigericin-induced NLRP3 inflammasome activation. **A, B** BMDMs prepared from C57BL/6 mice bone marrow were infected by lentivirus that expressing sh-Paxillin for 3 days. LPS-primed BMDMs were stimulated by MDP (50 μg/ml) for 6 h, ATP (5 mM) for 30 min, poly(dA:dT) (5 μg/ml) for 6 h, or Salmonella for 4 h. **C, D** LPS-primed BMDMs were stimulated by ATP (5 mM) for 30 min, Nigericin (2 μM) for 2 h, MSU (100 μg/ml) for 6 h, or Alum crystals (200 mg/ml) for 6 h. **E, F** BMDCs prepared from C57BL/6 mice bone marrow were infected by lentivirus that express sh-Paxillin for 3 days. LPS-primed BMDCs were stimulated by ATP (5 mM) for 30 min, Nigericin (2 μM) for 2 h, or MSU (100 μg/ml) for 6 h. IL-1β levels in supernatants were determined by ELISA (A, C, and E). Mature IL-1β (p17) and cleaved Casp-1 (p10) in supernatants as well as Paxillin, pro-IL-1β, and pro-Casp-1 in lysates were determined by Western blot (B, D, and F). **G, H** PBMCs isolated from healthy individuals were infected by lentivirus that expresses sh-Paxillin for 2 days. Before stimulation, PBMCs were treated with LPS (1 μg/ml) for 6 h, and PBMCs were then stimulated by ATP (5 mM) for 30 min, Nigericin (2 μM) for 2 h, MSU (100 μg/ml) for 6 h, Alum crystals (200 mg/ml) for 6 h, or MDP (50 μg/ml) for 6 h. Paxillin in lysates was determined by Western blot (G). IL-1β levels in supernatants were determined by ELISA (H). **I, J** THP-1 cells stably expressing sh-NC or sh-Paxillin were generated and differentiated to macrophages, which were then treated with ATP (5 mM) for 2 h, Nigericin (2 μM) for 2 h, or MSU (100 μg/ml) for 6 h. IL-1β levels in supernatants were determined by ELISA (I). Mature IL-1β (p17) and cleaved Casp-1 (p22/p20) in supernatants and Paxillin, pro-IL-1β and pro-Casp-1 in lysates were determined by Western blot (J). Data shown are means ± SEMs; ***p < 0.0001. ns, no significance.

Stable BMDMs were then treated with ATP, Nigericin, monosodium urate (MSU), and alum (Al). The results showed that ATP- and Nigericin-induced IL-1β secretion, IL-1β maturation, and Caspase-1 cleavage were attenuated by sh-Paxillin, whereas MSU- or Al-mediated IL-1β secretion, IL-1β maturation, and Caspase-1 cleavage were not affected by sh-Paxillin (Fig 7C, D). In addition, mouse bone marrow dendritic cells (BMDCs) stably expressed sh-Paxillin were constructed. Stable BMDCs were treated with ATP, Nigericin, and MSU. Similarly, ATP- and Nigericin-induced IL-1β secretion, IL-1β maturation, and Caspase-1 cleavage were reduced by sh-Paxillin, but MSU-mediated IL-1β secretion, IL-1β maturation, and Casp-1 cleavage were not affected by sh-Paxillin (Fig 7E, F). Moreover, human peripheral blood mononuclear cells (PBMCs) stably expressed sh-Paxillin were generated. The stable PBMCs were treated with ATP, Nigericin, MSU, Al, or MDP. ATP- and Nigericin-induced IL-1β secretion was down-regulated by sh-Paxillin, but MSU-, Al-, or MDP-mediated IL-1β secretion was not affected by sh-Paxillin, and the level of Paxillin was attenuated by sh-Paxillin (Fig 7G, H). Finally, THP-1 cells stably expressed sh-Paxillin were generated. The stable THP-1 cells were differentiated into macrophages, which were then treated with ATP, Nigericin, or MSU. ATP- and Nigericin-induced IL-1β secretion, IL-1β maturation and Caspase-1 cleavage were attenuated by sh-Paxillin, but MSU-mediated IL-1β secretion, IL-1β maturation, and Casp-1 cleavage were not affected by sh-Paxillin (Fig 7I, J). Collectively, these results demonstrate that Paxillin is essential for ATP- and Nigericin-induced NLRP3 inflammasome activation. Taken together, we reveal that Paxillin plays key roles in ATP-induced activation of P2X7 receptor and NLRP3 inflammasome by formatting the P2X7-Paxillin-NLRP3 complex (Fig 8).

**Figure 8.**
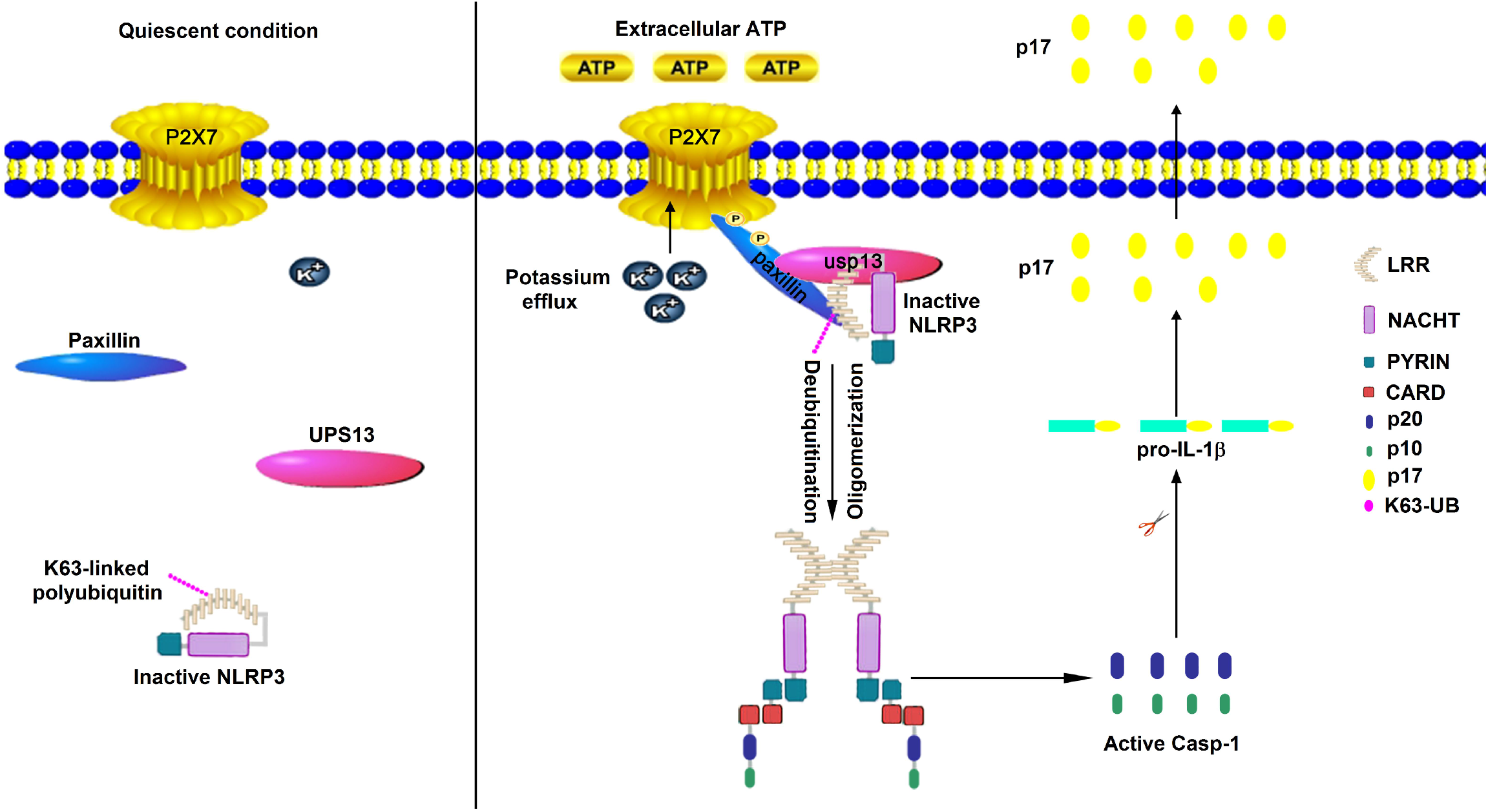
Paxillin regulates ATP-induced activation of P2X7 receptor and NLRP3 inflammasome by formatting the P2X7-Paxillin-NLRP3 complex. In the quiescent condition, the inactive, paxillin and usp13 distribute in the cytoplasm (left). However, in response to extracellular ATP, Paxillin is a molecular adaptor protein, which recruit uspl3 and NLRP3 at plasma membrane for the activation of NLRP3 inflammasome for efficient processing of P2X7.

## Discussion

NLRP3 inflammasome activation requires P2X7 receptor stimulation mediated by ATP (Di Virgilio et al., 2017; Latz et al., 2013; Surprenant and North, 2009). Upon ATP treatment, P2X7 induces transmembrane K^+^ ion efflux and Pannex-1 hemichannel to form large pore for inflammasome activation (Pelegrin and Surprenant, 2006; Pelegrin and Surprenant, 2007). However, the molecule connecting P2X7 receptor and NLRP3 inflammasome has not been revealed. This study identifies that Paxillin plays key roles in ATP-induced activation of P2X7 receptor and NLRP3 inflammasome by promoting the formation of P2X7-Paxillin-NLRP3 complex. The primary function of Paxillin is a molecular adapter or scaffold protein that provides multiple docking sites at the plasma membrane for processing of Integrin- and growth factor-mediated signals (Ishibe et al., 2003). Initially, we revealed a direct interaction between Paxillin and NLRP3. The function and localization of Paxillin is tightly modulated by phosphorylation (Webb et al., 2005). Phosphorylations at Y31 and Y118 sites in FAK- and Src-dependent manners (Schaller and Parsons, 1995) are essential for interaction between Paxillin and downstream effectors, such as ERK (Hanks et al., 2003) and Crk2 (Valles et al., 2004). Tyrosine phosphorylation of Paxillin regulates the assembly and turnover of adhesion complexes (Lopez-Colome et al., 2017). Here, we demonstrate that ATP-induced Paxillin Y118 phosphorylation is essential for Paxillin-NLRP3 interaction and Paxillin promotes NLRP3 inflammasome activation upon ATP treatment.

Extracellular ATP stimulates the P2X7 ATP-gated ion channel, triggers K^+^ efflux, and induces gradual recruitment of the Pannexin-1 membrane pore, leading to inflammasome activation (Kahlenberg and Dubyak, 2004; Kanneganti et al., 2007; Mariathasan et al., 2006). Interestingly, Paxillin interacts with P2X7, ATP promotes P2X7-Paxillin interaction, Paxillin Y31 phosphorylation is required for such interaction, P2X7 K30 is essential for P2X7-Paxillin association. Thus, we reveal that ATP stimulates P2X7-Paxillin interaction, induces K^+^ efflux, activates Paxillin phosphorylation, and promotes Paxillin-NLRP3 interaction, and suggest that Paxillin plays a key role in the P2X7-Paxillin-NLRP3 complex assembly. Paxillin is a downstream adaptor of Integrin and functions as a scaffold for protein recruitment into a complex (Franceschini et al., 2015). Our finding reveals a new function of Paxillin in the recruitment of proteins involved in ATP-induced activation of P2X7 receptor and NLRP3 inflammasome.

The molecular mechanism underlying the function of Paxillin in activation of P2X7 receptor and NLRP3 inflammasome is determined. As NLRP3 deubiquitination is important for the inflammasome activation (Py et al., 2013), the roles of ATP and Paxillin in NLRP3 deubiquitination were evaluated. Notably, Paxillin facilitates the remove of NLRP3 K63-linked ubiquitination at K689, and ATP as well as K^+^ efflux are critical for this regulation. The importance of K^+^ efflux in NLRP3 inflammasome activation has been revealed (Piccini et al., 2008). The drop of K^+^ concentration triggers NLRP3 inflammasome, whereas a high concentration of K^+^ blocks NLRP3 inflammasome (Munoz-Planillo et al., 2013). However, the molecular mechanism between decreased levels of intracellular K^+^ and NLRP3 activation remains largely unknown. Recent study has showed that intracellular K^+^ reduction mediated by P2X7 enhances NLRP3 interaction with NEK7 (He et al., 2016). The present study identifies that Paxillin is essential for ATP-induced NLRP3 inflammasome activation and for NLRP3 recruitment in the plasma membrane.

Post-translational modifications (PTMs), including ubiquitination (Py et al., 2013), phosphorylation (Song and Li, 2018), and Sumoylation (Barry et al., 2018) are critical for inflammasome activation. NLRP3 deubiquitination is required for NLRP3 inflammasome activation and pharmacological inhibition of NLRP3 deubiquitination impairs the inflammasome activation (Py et al., 2013). ATP and Nigericin induce NLRP3 deubiquitination and inflammasome activation (Juliana et al., 2012). We demonstrate that ATP-induced Paxillin Y118 phosphorylation promotes the remove of NLRP3 K63-linked ubiquitination at K689. Additionally, we discovered that UPS13 is the deubiquitinating enzyme required for Paxillin-mediated NLRP3 deubiquitination. UPS13 interacts with Paxillin and NLRP3, phosphorylation of Paxillin at Y31 is important for Paxillin-UPS13 interaction, and UPS13 is essential for Paxillin-mediated NLRP3 deubiquitination upon ATP treatment.

ATP induces Paxillin-containing membrane protrusions (Silber et al., 2012) and Paxillin acts as a scaffold for protein recruitments into a complex apposing to the plasma membrane (Franceschini et al., 2015). Confocal microscopy analysis show that ATP induces the translocation of endogenous NLRP3 and endogenous Paxillin from the cytosol to the plasma membrane. Membrane flotation assay indicate that NLRP3, Paxillin, and P2X7 colocalize to the plasma membrane upon ATP treatment. Therefore, we suggest that ATP induces Paxillin and NLRP3 membrane migration to facilitate P2X7-Paxillin-NLRP3 complex formation.

The biological role of Paxillin in NLRP3 inflammasome activation is determined. We reveal that Paxillin is essential for ATP-induced NLRP3 inflammasome activation, including IL-1β secretion, IL-1β maturation, and Caspase-1 cleavage. After the response to tissue damage and cellular stress, ATP released to enhance tissue repair and promote the recruitment of immune phagocytes and dendritic cells (Stagg and Smyth, 2010). ATP-P2X7 receptor signaling is involved in many pathological conditions, including infectious diseases, inflammatory diseases, and neurodegenerative disorders (Savio et al., 2018). We demonstrate that Paxillin is a molecular adaptor that recruits UPS13 and NLRP3 on plasma membrane for NLRP3 inflammasome activation, and thereby may act as a potential target of therapeutics to inflammatory diseases.

## Materials and Methods

### Animal study

Mouse BMDCs were differentiated from fresh bone marrow cells of C57BL/6 WT mice in RPMI 1640 medium containing 10% heat-inactive fetal bovine serum (FBS) in the presence of granulocyte macrophage colony-stimulating factor (GM-CSF) in six-well plates for 6 days. The culture medium was replaced every other day.

Mouse BMDMs were differentiated from fresh bone marrow cells of C57BL/6 WT mice. The bone marrow cells were incubated in six-well plates for 6 days with 10% L929-conditioned, 10% heat-inactive FBS in RPMI 1640 medium. The culture medium was replaced every 2 days. The animal study was approved by the Institutional Review Board of the College of Life Sciences, Wuhan University, and was conducted in accordance with the guidelines for the protection of animal subjects.

### Cell lines and cultures

Hela and human embryonic kidney cells (HEK 293T) were purchased from American Type Culture Collection (ATCC) (Manassas, VA, USA). Human acute monocytic leukemia cell line (THP-1) was a gift from Dr. Jun Cui of State Key Laboratory of Biocontrol, School of Life Sciences, Sun Yat-sen University, Guangzhou 510275, PRC.THP-1 cells were cultured in RPMI 1640 medium supplemented with 10% heat-inactivated FBS, 100 U/ml penicillin, and 100 μg/ml streptomycin sulfate. Hela and HEK293T cells were cultured in Dulbecco modified Eagle medium (DMEM) purchased from Gibco (Grand Island, NY, USA) supplemented with 10% FBS, 100 U/ml penicillin, and 100 μg/ml streptomycin sulfate. Hela, HEK293T and THP-1 cells were maintained in an incubator at 37°C in a humidified atmosphere of 5% CO_2_.

### Reagents

phorbol-12-myristate-13-acetate (TPA) (P1585), OptiPrepTM (D1556) and DMSO (D8418) were purchased from Sigma-Aldrich (St. Louis, MO, USA). RPMI 1640 and Dulbecco modified Eagle medium (DMEM) were obtained from Gibco (Grand Island, NY, USA). Lipopolysaccharide (LPS) (tlrl-b5lps), adenosine triphosphate (ATP) (987-65-5), Nigericin (28643-80-3), dA:dT(86828-69-5), MDP (53678-77-6), Glybenclamide (10238-21-8), MSU (1198-77-2), Alum Crystals (7784-24-9) and Ac-YVAD-cmk (178603-78-6) were obtained from InvivoGene Biotech Co., Ltd. (San Diego, CA, USA). A438079 (899431-18-6), AZ10606120 (607378-18-7) and BAPTA-AM (126150-97-8) were purchased form Tocris Bioscience. Antibody against Flag (F3165), HA (H6908) and monoclonal mouse anti-GAPDH (G9295) were purchased from Sigma (St. Louis, MO, USA). Monoclonal rabbit anti-NLRP3 (D2P5E), Ubiquitin mouse mAb (P4D1), Monoclonal rabbit anti-K63-linkage Specific Polyubiquitin (D7A11), Monoclonal rabbit anti-caspase-3 (13809S), Monoclonal rabbit anti-P2X7 (9662S), Monoclonal rabbit anti-p-paxillin(Tyr118) (2541S), Monoclonal rabbit anti-calnexin (C5C9), monoclonal rabbit anti-IL-1β (D3U3E), IL-1β mouse mAb (3A6) and monoclonal rabbit anti-caspase-1 (catalog no. 2225) were purchased from Cell Signaling Technology (Beverly, MA, USA). Monoclonal mouse anti-ASC (sc-271054), polyclonal rabbit anti-caspase-1 p10 (sc-515) and polyclonal rabbit anti-IL-1β (sc-7884) were purchased from Santa Cruz Biotechnology (Santa Cruz, CA, USA). Polyclonal Goat anti-mouse IL-1β (p17) (AF-401-NA) was from R&D Systems. Monoclonal mouse anti-paxillin (7019741) was purchased from BD Biosciences. Polyclonal rabbit anti-p-paxillin (Tyr31) (1690606A) was purchased from life technologiex. Monoclonal mouse anti-NLRP3 (AG-20B-0014-C100) was purchased from adipogen to detection endogenous NLRP3 in THP1 cells, BMDC and BMDM. Lipofectamine 2000, normal rabbit IgG and normal mouse IgG were purchased from Invitrogen Corporation (Carlsbad, CA, USA).

### Plasmid construction

The cDNAs encoding human paxillin, NLRP3, ASC, pro-Casp-1 and IL-1β were obtained by reverse transcription of total RNA from TPA-differentiated THP-1 cells, followed by PCR using specific primers. The cDNAs were sub-cloned into pcDNA3.1(+) and pcagg-HA vector. The pcDNA3.1(+)-3×Flag vector was constructed from pcDNA3.1(+) vector through inserting the 3×Flag sequence between the NheI and HindIII site. Following are the primers used in this study.

Flag-NLRP3: 5’-CGCGGATCCATGAAGATGGCAAGCACCCGC-3’, 5’-CCGCTCGAGCTACCAAGAAGGCTCAAAGAC-3’; Flag-ASC: 5’-CCGGAATTCATGGGGCGCGCGCGCGACGCCAT-3’, 5’-CCGCTCGAGTCAGCTCCGCTCCAGGTCCTCCA-3’; Flag-Casp-1: 5’-CGCGGATCCATGGCCGACAAGGTCCTGAAG-3’, 5’-CCGCTCGAGTTAATGTCCTGGGAAGAGGTA-3’; Flag-paxillin: 5’-CCGGAATTCTATGGACGACCTCGACGCCCT-3’, 5’-CCGCTCGAGCTAGCAGAAGAGCTTGAGGAA-3’; Myc-paxillin: 5’-CCGGAATTCTATGGACGACCTCGAC-3’, 5’-CCGCTCGAGCTAGCAGAAGAGCTTG-3’; Myc-paxillin(y118a): 5’-GTGAGGAGGAGCACGTCGCAAGCTTCCCCAACAAGCAGAA-3’, 5’-ATTTCTGCTTGTTGGGGAAGCTTGCGACGTGCTCCTCCTC-3’; HA-paxillin: 5’-CCGGAATTCATGGACGACCTCGACGCCCTG-3’, 5’-CGCTCGAGGCAGAAGAGCTTGAGGAAGCA-3’; HA-paxillin(y31a): 5’-GCCTGTGTTCTTGTCGGAGGAGACCCCCGCATCATACCCA-3’, 5’-GTGTGGTTTCCAGTTGGGTATGATGCGGGGGTCTCCTCCG-3’; HA-paxillin(y118a): 5’-GTGAGGAGGAGCACGTCGCAAGCTTCCCCAACAAGCAGAA-3’, 5’-ATTTCTGCTTGTTGGGGAAGCTTGCGACGTGCTCCTCCTC-3’; HA-NLRP3: 5’-TACGAGCTCATGAAGATGGCAAGCACCCGC-3’, 5’-CCGCTCGAGCCAAGAAGGCTCAAAGACGAC-3’; pGEX-6p-1-paxillin: 5’-CCGGAATTCATGGACGACCTCGACGCCCTG-3’, 5’-CGCTCGAGGCAGAAGAGCTTGAGGAAGCA-3’; pGEX-6p-1-LRR: 5’-CGCGGATCCATGTCTCAGCAAATCAGGCTG-3’, 5’-CCGCTCGAGCTACCAAGAAGGCTCAAAGAC-3’; AD-paxillin: 5’-CCGGAATTCATGGACGACCTCGACGCCCTG-3’, 5’-CGCTCGAGGCAGAAGAGCTTGAGGAAGCA-3’; pGBKT7-LRR: 5’-CGCGGATCCATATGTCTCAGCAAATCAGGC-3’, 5’-CCGCTCGAGCTACCAAGAAGGCTCAAAGAC-3’;

The P2X7 and usp13 truncates was cloned into pcDNA3.1(+) and the PYRIN, NACHT, and LRR domain of NLRP3 protein was cloned into pcDNA3.1(+) and pcaggs-HA vector using specific primers. Following are the primers used in this study. Flag-PYRIN: 5’-AAAGGATCCATGAAGATGGCAAGCACCCGC-3’, 5’-CGGCTCGAGCTATAAACCCATCCACTCCTCTTC-3’; Flag-NACHT: 5’-AAAGGATCCCTGGAGTACCTTTCGAGAATCTC-3’, 5’-CCCCTCGAGCTAGATCTTGCAACTTAATTTCTTC-3’; Flag-LRR: 5’-AAAGGATCCTCTCAGCAAATCAGGCTGGAG-3’, 5’-CGGCTCGAGCTACCAAGAAGGCTCAAAGACG-3’; Flag-P2X7: 5’-ATTGGTACCATGCCGGCCTGCTGCAGCTGCAGT-3’, 5’-CCGCTCGAGTCAGTAAGGACTCTTGAAGCCACT-3’; Flag-P2X7(26-595): 5’-CAAGATATCATGTATGGCACCATTAAGTGG-3’, 5’-CCGCTCGAGTCAGTAAGGACTCTTGAAGCC-3’; Flag-P2X7(47-595): 5’-AATGATATCATGAGTGACAAGCTGTACCAG-3’, 5’-CCGCTCGAGTCAGTAAGGACTCTTGAAGCC-3’; Flag-P2X7(335-595): 5’-CCGGAATTCTATGGTGTACATCGGCTCAAC-3’, 5’-CCGCTCGAGTCAGTAAGGACTCTTGAAGCC-3’; Flag-P2X7(356-595): 5’-CCGGAATTCTATGGACACTTACTCCAGTAA-3’, 5’-CCGCTCGAGTCAGTAAGGACTCTTGAAGCC-3’; Flag-P2X7(27-595): 5’-CGGGGTACCATGGGCACCATTAAGTGGTTC-3’, 5’-CTAGTCTAGATCAGTAAGGACTCTTGAAGC-3’; Flag-P2X7(29-595): 5’-CGGGGTACCATGATTAAGTGGTTCTTCCAC-3’, 5’-CTAGTCTAGATCAGTAAGGACTCTTGAAGC-3’; Flag-P2X7(30-595): 5’-CGGGGTACCATGAAGTGGTTCTTCCACGTG-3’, 5’-CTAGTCTAGATCAGTAAGGACTCTTGAAGC-3’; Flag-P2X7(31-595): 5’-CGGGGTACCATGTGGTTCTTCCACGTGATC-3’, 5’-CTAGTCTAGATCAGTAAGGACTCTTGAAGC-3’; Flag-P2X7(32-595): 5’-CGGGGTACCATGTTCTTCCACGTGATCATC-3’, 5’-CTAGTCTAGATCAGTAAGGACTCTTGAAGC-3’; Flag-P2X7(33-595): 5’-CGGGGTACCATGTTCCACGTGATCATCTTT-3’, 5’-CTAGTCTAGATCAGTAAGGACTCTTGAAGC-3’; Flag-P2X7(34-595): 5’-CGGGGTACCATGCACGTGATCATCTTTTCC-3’, 5’-CTAGTCTAGATCAGTAAGGACTCTTGAAGC-3’; Flag-P2X7(35-595): 5’-CGGGGTACCATGGTGATCATCTTTTCCTAC-3’, 5’-CTAGTCTAGATCAGTAAGGACTCTTGAAGC-3’; Flag-P2X7(36-595): 5’-CGGGGTACCATGATCATCTTTTCCTACGTT-3’, 5’-CTAGTCTAGATCAGTAAGGACTCTTGAAGC-3’; Flag-P2X7(k30a): 5’-AATTATGGCACCATTGCGTGGTTCTTCCACGTGAT-3’, 5’-ATGATCACGTGGAAGAACCACGCAATGGTGCCATA-3’; Flag-USP13: 5’-CGGGGTACCATGCAGCGCCGGGGCGCCCTG-3’, 5’-CCGCTCGAGTTAGCTTGGTATCCTGCGGTA-3’; Flag-usp13(1-300): 5’-CGGGGTACCATGCAGCGCCGGGGCGCCCTG-3’, 5’-CCGCTCGAGTTACCCATGCATATGAAGCAT-3’; Flag-usp13(301-624): 5’-CGGGGTACCATGACAGAGAATGGGCTCCAG-3’, 5’-CCGCTCGAGTTATTCCTCTCCTGGCTGTAA-3’; Flag-usp13(301-863): 5’-CGGGGTACCATGACAGAGAATGGGCTCCAG-3’, 5’-CCGCTCGAGTTAGCTTGGTATCCTGCGGTA-3’; Flag-usp13(625-863): 5’-CGGGGTACCATGGAACTTCCAGACATCAGC-3’, 5’-CCGCTCGAGTTAGCTTGGTATCCTGCGGTA-3’. HA-PYRIN: 5’-CCGGAATTCATGAAGATGGCAAGCACCCGC-3’, 5’-CCGCTCGAGTAAACCCATCCACTCCTCTTC-3’; HA-NACHT: 5’-CCGGAATTCATGCTGGAGTACCTTTCGAGA-3’, 5’-CCGCTCGAGGATCTTGCAACTTAATTTCTT-3’; HA-LRR: 5’-ATCGAGCTCATGTCTCAGCAAATCAGGCTG-3’, 5’-CCGCTCGAGCCAAGAAGGCTCAAAGACGAC-3’;

### Lentivirus Production and Infection

The targeting sequences of shRNAs for the human Paxillin and mouse paxillin were as follows: Human sh-Paxillin: 5’-CCCGACCTAATTGTCTTTGTT-3’; Mouse sh-Paxillin: 5’-TCTGAACTTGACCGGCTGTTA-3’. A PLKO.1 vector encoding shRNA for a negative control (Sigma-Aldrich, St. Louis, MO, USA) or a specific target molecule (Sigma-Aldrich) was transfected into HEK293T cells together with psPAX2 and pMD2.G with Lipofectamine 2000. Culture supernatants were harvested 36 and 60 h after transfection and then centrifuged at 2,200 rpm for 15 min. THP-1 cells were infected with the supernatants contain lentiviral particles in the presence of 4 μg/ml polybrene (Sigma). After 48 h of culture, cells were selected by 1.5 μg/ml puromycin (Sigma) for 5days. The BMDM and BMDCs cells were infected with lentiviral particles for 1day and treated with 1.5 μg/ml puromycin (Sigma) for 2 days. The results of each sh-RNA-targeted protein were detected by immunoblot analysis.

We using the 3*Flag sequence to replace the GFP protein in the pLenti CMV GFP Puro vector (Addgene, 658-5) for adding some Restriction Enzyme cutting site (XbaI-EcoRV-BstBI-BamHI) before the 3 ×Flag tag. Then the pLenti vector encoding paxillin protein was transfected into HEK293T cells together with psPAX2 and pMD2.G with Lipofectamine 2000. Following are the primers used in this study. pLenti-paxillin: 5’-CTAGTCTAGAATGGACGACCTCGACGCCCT-3’, 5’-GATTTCGAAGCAGAAGAGCTTGAGGAAGCA-3’. Culture supernatants were harvested 36 and 60 h after transfection and then centrifuged at 2,200 rpm for 15 min. THP-1 cells were infected with the supernatants contain lentiviral particles in the presence of 4 μg/ml polybrene (Sigma). After 48 h of culture, cells were selected by 1.5 μg/ml puromycin (Sigma) for 5 days. The results of paxillin protein was detected by immunoblot analysis.

### Enzyme-linked immunosorbent assay (ELISA)

The concentrations of human IL-1β in culture supernatants were measured by ELISA kit (BD Biosciences, San Jose, CA, USA). The mouse IL-1β ELISA Kit was purchased from R&D.

### THP-1 macrophages stimulation

THP-1 cells were differentiated to macrophages with 60 nM phorbol-12-myristate-13-acetate (TPA) for 12–14 h, and cells were cultured for 24 h without TPA. And then the differentiated cells were stimulated in 6 cm plates with Nigericin or ATP. Supernatants were collected for measurement of IL-1β by ELISA. Cells were harvested for immunoblot analysis.

### Activated caspase-1 and mature IL-1β measurement

The supernatant of the cultured cells was collected for 1 ml in the cryogenic vials (Corning). The supernatant was frozen in −80°C for 4 h. The Rotational Vacuum concentrator machine which was purchase from Martin Christ was used for the freeze drying. The drying product was dissolved in 100 μl PBS and mixed with SDS loading buffer for western blotting analysis with antibodies for detection of activated caspase-1 (D5782 1:500, Cell Signaling), mature IL-1β (Asp116 1:500, Cell Signaling), polyclonal rabbit anti-caspase-1 p10 (sc-515) or Polyclonal Goat anti-mouse IL-1β (p17) (AF-401-NA). Adherent cells in each well were lysed with the lyses buffer described below, followed by immunoblot analysis to determine the cellular content of various protein.

### Western blot analysis

HEK293T whole-cell lysates were prepared by lysing cells with buffer (50 mM Tris-HCl, pH7.5, 300 mM Nacl, 1% Triton-X, 5 mM EDTA and 10% glycerol). The TPA-differentiated THP-1 cells lysates were prepared by lysing cells with buffer (50 mM Tris-HCl, pH7.5, 150 mM NaCl, 0.1% Nonidetp 40, 5 mM EDTA and 10% glycerol). Protein concentration was determined by Bradford assay (Bio-Rad, Hercules, CA, USA). Cultured cell lysates (30 μg) were electrophoresed in an 8–12% SDS-PAGE gel and transferred to a PVDF membrane (Millipore, MA, USA). PVDF membranes were blocked with 5% skim milk in phosphate buffered saline with 0.1% Tween 20 (PBST) before being incubated with the antibody. Protein band were detected using a Luminescent image Analyzer (Fujifilm LAS-4000).

### Co-immunoprecipitation assays

HEK293T whole-cell lysates were prepared by lysing cells with buffer (50 mM Tris-HCl, pH7.5, 300 mM NaCl, 1% Triton-X, 5 mM EDTA and 10% glycerol). TPA-differentiated THP-1 cells lysates were prepared by lysing cells with buffer (50 mM Tris-HCl, pH7.5, 150 mM NaCl, 0.1% Nonidetp40, 5 mM EDTA and 10% glycerol). Lysates were immunoprecipitated with control mouse immunoglobulin G (IgG) (Invitrogen) or anti-Flag antibody (Sigma, F3165) with Protein-G Sepharose (GE Healthcare, Milwaukee, WI, USA).

### Confocal microscopy

HEK293T cells and Hela cells were transfected with plasmids for 24–36 h. Cells were fixed in 4% paraformaldehyde at room temperature for 15 min. After being washed three times with PBS, permeabilized with PBS containing 0.1% Triton X-100 for 5 min, washed three times with PBS, and finally blocked with PBS containing 5% BSA for 1 h. The cells were then incubated with the monoclonal mouse anti-Flag antibody (F3165, Sigma) and Monoclonal rabbit anti-HA (H6908, Sigma) overnight at 4°C, followed by incubation with FITC-conjugate donkey anti-mouse IgG (Abbkine) and Dylight 649-conjugate donkey anti-rabbit IgG (Abbkine) for 1 h. After washing three times, cells were incubated with DAPI solution for 5 min, and then washed three more times with PBS. Finally, the cells were analyzed using a confocal laser scanning microscope (Fluo View FV1000; Olympus, Tokyo, Japan).

### GST pull down assays

The plasmids pGEX6p-1-paxillin and pGEX6p-1-LRR were transfected into Escherichia coli strain BL21. After growing in LB medium at 37°C until the OD600 reached 0.6–0.8, Isopropyl β-D-1-thiogalactopyranoside (IPTG) was added to a final concentration of 1 mM and the cultures grew for an additional 4 h at 37°C for GST-Paxillin and GST-LRR protein. And then the GST protein, GST-paxillin protein and GST-LRR protein were purified from E. coli bacteria. For GST-Paxillin pull-down assay, glutathione-Sepharose beads (Novagen) were incubated with GST-Paxillin or GST protein. After washed with phosphate-buffered saline (PBS), these beads were incubated with cell lysates from HEK293T which were transfected with plasmids encoding Flag-NLRP3 for 4 h at 4°C. The precipitates were washed three times, boiled in 2 × SDS loading buffer, separated by 10% SDS-PAGE, immunoblotted with anti-GST, anti-Flag. It was the same for the GST-LRR pull down assay.

### Yeast two-hybrid analyses

Saccharomyces cerevisiae strain AH109, control vectors pGADT7, pGBKT7, pGADT7-T, pGBKT7-Lam and pGBKT7-p53 were purchased from Clontech (Mountain View, CA, USA). Yeast strain AH109 was co-transformed with the combination of the pGADT7 and the pGBKT7 plasmids. Transformed yeast cells containing both plasmids were first grown on SD-minus Trp/Leu plates (DDO) to maintain the two plasmids and then were sub-cloned replica plated on SD-minus Trp/Leu/Ade/His plate (QDO).

### Subcellular Fractionation

To separate cell membranes from the soluble cellular components, cells were washed once in hypotonic buffer (10 mM Tris-HCl, pH7.4, 10 mM KCl, 1.5 mM MgCl) supplemented with a protease inhibitor cocktail (Roche), incubated on ice in hypotonic buffer, and lysed by dounce homogenization. Lysates were centrifuged at 4°C for 5 minutes at 2,500 x g to get the supernatants. Supernatants were centrifuged at 100,000 x g for 1 h at 4°C. The resultant pellets (membrane fraction) were resuspended in lysis buffer 1/5 volumes equal to those of the supernatants (cytosolic fraction), stored with the addition of 6 x Laemmli Buffer, and analyzed by Western blot.

For membrane floatation assays, post nuclear supernatants were collected and described above, mixed with OptiprepTM-supplemented hypotonic buffer to yield a final concentration of 45% optiprep at laid at the bottom of an OptiprepTM step gradient ranging from 10% (top) to 45% (bottom), and spun at 52,000 x g for 90 minutes. Gradient was then fractionated into 24 fractions and analyzed by western blot. For gradients run in the presence of Triton X-100, post nuclear supernatants were mixed with a 10% Triton X-100 solution to achieve a final concentration of 1%.

### Statistical analyses

All experiments were reproducible and repeated at least three times with similar results. Parallel samples were analyzed for normal distribution using Kolmogorov-Smirnov tests. Abnormal values were eliminated using a follow-up Grubbs test. Levene’s test for equality of variances was performed, which provided information for Student’s t-tests to distinguish the equality of means. Means were illustrated using histograms with error bars representing the SD; a P value of <0.05 was considered statistically significant.

## Acknowledgments

This work was supported by National Natural Science Foundation of China (81730061, 81902053 and 81471942), National Health and Family Planning Commission of China (National Mega Project on Major Infectious Disease Prevention) (2017ZX10103005 and 2017ZX10202201), and China Postdoctoral Science Foundation (2018T110923).

## Author Contributions

W.W., D.H., K.W., Y.L., and J.W. contributed to the design of experiments. W.W., D.H., Y.F., C.W., A.L., W.L., Y.W., K.C., M.T., F.X., Q.Z., W.C., P.P., and P.W. contributed to the conduction of experiments. W.W., D.H., Y.F., W.L., Q.Z., C.W., A.L., W.L., P.P., P.W., K.W., Y.L., and J.W. contributed to the reagents. W.W., D.H., Y.F., and J.W. contributed to the writing the paper. W.W., and J.W. contributed to the editing the paper.

